# Endothelial Dnmt3a controls placenta vascularization and function to support fetal growth

**DOI:** 10.1101/2022.07.28.501807

**Authors:** Stephanie Gehrs, Moritz Jakab, Ewgenija Gutjahr, Zuguang Gu, Dieter Weichenhan, Carolin Mogler, Matthias Schlesner, Christoph Plass, Hellmut G. Augustin, Katharina Schlereth

## Abstract

The fetoplacental capillary network is of vital importance for proper nourishment during early development. Inadequate maternal-fetal circulation has emerged as one of the main pathophysiological features of placental insufficiency. Meta-analysis of human placental endothelial cells (EC) revealed that downregulation of the *de novo* DNA methyltransferase 3A (*DNMT3A*) is associated with preeclampsia. However, mechanistic insights into functional consequences of altered DNA methylation patterns during placental vascular development remain elusive. Here, we investigated the role of *Dnmt3a* in the vasculature during murine placenta development. Spatial and temporal expression analyses revealed an induction of *Dnmt3a* in the mature labyrinth layer. The global and endothelium-specific loss (ECKO) of *Dnmt3a* resulted in reduced placental vascularization and fetal growth restriction. EC deleted for *Dnmt3a* demonstrated extensive loss of DNA methylation, particularly close to angiogenesis related genes. Loss of DNA methylation decreased the angiogenic capacity of EC *in vitro* and *in vivo*. Collectively, these data identify DNMT3A as the main DNA methyltransferase in the human and murine placental vasculature and characterize its importance for physiological endothelial function. The Dnmt3a-dependent regulation of genes related to placenta insufficiency validates Dnmt3a ECKO mice as an epigenetically driven mouse model of placenta insufficiency with potential translational relevance.

## Introduction

The placenta represents a temporary organ that acts at the interface between mother and fetus. Besides its roles in the production of pregnancy-supporting hormones and building a selective barrier to fight against internal infection, the supply of nutrients and oxygen to the developing embryo represents the most vital task of the placenta (Rossant and Cross 2001). In the human placenta, the exchange of nutrients and oxygen between the maternal and the fetoplacental circulation takes place in the chorionic villi. The maternal-fetal interface transports oxygen and nutrients from the maternal bloodstream to the complex branched fetoplacental capillary network representing the functional unit of the placenta. The human chorionic villi and the mouse labyrinth represent analogous structures based on the morphological and functional similarity (Georgiades et al. 2002; Woods et al. 2018).

Abnormal placental development affects the placental architecture and impairs placental function resulting in impaired fetal growth. Preeclampsia (PE) and intrauterine growth restriction (IUGR; syn. fetal growth restriction, FGR) are severe pregnancy disorders that share common pathologies. PE is the main cause of maternal and perinatal morbidity affecting 2-8% of pregnancies worldwide in the early third trimester (Duley 2009; Rana et al. 2019; Steegers et al. 2010). This disease is characterized by inadequate trophoblast invasion into the uterine wall and defective remodeling of the placental vasculature leading to vascular dysfunction (Opichka et al. 2021). FGR affects approximately 5 - 8% of all pregnancies and refers to a condition in which the fetus is unable to achieve its genetically predetermined growth potential (Romo et al. 2009). A major cause of FGR is placental insufficiency resulting in inadequate fetal supply and hence, reduced fetal growth. However, the exact pathomechanism of these pregnancy disorders is still largely unknown.

Given the pivotal role of the maternal-fetal circulation for fetal nourishment, its dysregulation has emerged as one of the main pathophysiological features of placental insufficiency. In fact, abnormal growth of the fetoplacental endothelium has been described as the underlying cause of fetal growth restriction (Kingdom et al. 2000; Mayhew et al. 2003; Teasdale 1984). Similarly, studies indicate that compromised placenta circulation and dysregulation of angiogenic molecules represent common features of PE and IUGR/FGR (Karumanchi et al. 2009; Rana et al. 2022). Yet, the underlying mechanisms responsible for dysfunctional placental circulation remain largely elusive.

Highly complex epigenetic mechanisms including DNA methylation and histone posttranslational modifications, among others, represent essential control mechanisms of tissue function and development by shaping gene expression (Jaenisch and Bird 2003). As DNA methylation patterns are inherited during cell division, it represents a stable marker for epigenetic alterations and is therefore central for epigenetic cell memory (Schübeler 2015). DNA methylation occurs primarily on the cytosines of CpG dinucleotides. In mammals, *de novo* establishment of DNA methylation is governed by DNA methyltransferase 3 (DNMT3) A and B (Okano et al. 1998). In mice, the global deletion of *Dnmt3a* leads to reduced animal size with premature death at four weeks of age (Okano et al. 1999). Yet, the mechanistic understanding of the impact of DNA methylation alterations on vascular function in the context of dysfunctional placentas is extremely limited.

Several studies suggest that DNA methylation alterations in the placenta are involved in the pathogenesis of severe placental pathologies (Cirkovic et al. 2020; Nelissen et al. 2011; Wilson et al. 2018). While global DNA hypermethylation in the placenta can be associated with being born large for gestational age, specific hypomethylation patterns are observed in IUGR placentas (Banister et al. 2011; Dwi Putra et al. 2020). Recent studies link DNMT3A to placenta insufficiency. For instance, downregulation of *DNMT3A* in PE placentas has been associated with reduced migration capacity of trophoblasts (Jia et al. 2017). While most research focuses on trophoblast epigenetics, one study indicates that reduction of DNMT3A is specifically detectable in the fetal capillaries of the chorionic villi of placentas from early growth arrest cases (Gu et al. 2017). These results demonstrate that abnormal placentation is linked to DNA methylation dysregulation.

Accordingly, identifying epigenetic aberrations in placenta insufficiency may help to develop epigenetic therapies against PE and IUGR. Considering the functional importance of the fetoplacental capillary network for fetal nourishment, the aims of this study were to assess the role of DNA methylation modifiers in human placenta endothelium during health and preeclampsia by a meta-analytic approach and to gain further mechanistic insight using mouse models. The comparative analysis of diseased vs. healthy and fetal vs. maternal EC retrieved from single-cell data sets (Tsang et al. 2017; Marsh and Blelloch 2020) revealed a prominent role of the DNA methylation modifier Dnmt3a in physiological fetoplacental vascularization. *In vivo*, loss of *Dnmt3a* led to DNA hypomethylation, reduced angiogenic capacity of EC and, consequently, to decreased angiogenesis in the placenta. Further analyses propose placenta insufficiency in *Dnmt3a* KO mice. Altogether, our study presents DNMT3A as a critical regulator of placenta vascularization and function in humans and mice.

## Results

### Reduced *DNMT3A* expression in human placenta endothelial cells is associated with preeclampsia

To examine the role of the endothelium in placenta function and preeclamptic dysfunction, we analyzed published single-cell RNA sequencing data of 1368 and 756 EC sampled from full-term and early preeclamptic human placentas, respectively (Tsang et al. 2017). Shared nearest neighbor (SNN) clustering of single-cells from healthy placenta tissue allowed to classify different cell populations based on congruent marker expression, including different types of trophoblasts lineages and EC (**Figure 1A**). To assess the importance of distinct epigenetic mechanisms in placenta cell types, the expression of genes involved in writing, reading or erasing epigenetic marks (chromatin remodeling, histone modification, DNA methylation modification; see Plass et al. 2013) was analyzed. The expression of DNA methylation modifiers (e.g. *DNMT1, GADD45B, MGMT*) was highest in Hofbauer cells, EC and trophoblasts, indicating an important role of DNA methylation in those cell types. Of the *de novo* DNA methyltransferases, *DNMT3A*, but not *DNMT3B* was highly expressed in placenta cells, particularly in trophoblasts and EC (**Fig. 1B**). These data suggest DNMT3A as a key epigenetic DNA methylation modifier in the placenta endothelium.

**Figure 1:**
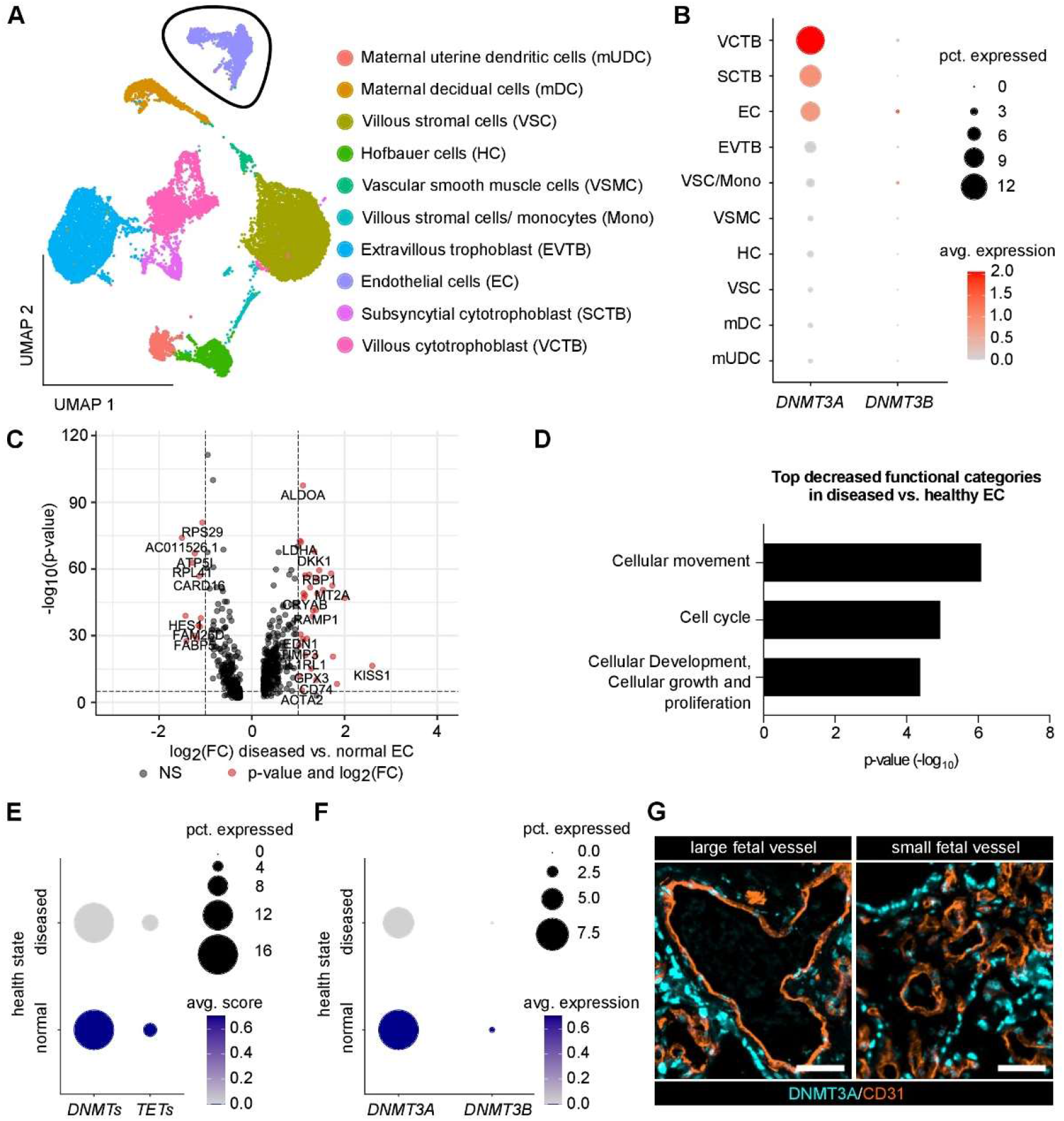
Reduced *DNMT3A* expression in human placenta EC is associated with preeclampsia. **A** UMAP of healthy human placenta single-cell data (Tsang et al., 2017) with cell type annotation based on marker gene expression. Endothelial cells are encircled in black. **B** *DNMT3A* and *DNMT3B* expression in different cell types of healthy human placenta. Cell types are sorted descending according to the percentage of cells expressing *DNMT3A*. **C** Volcano plot visualizing the gene expression changes (717 genes total) between diseased (= preeclampsia) and healthy placenta EC (red dots indicate significantly differentially expressed genes). Cutoff for the p-value (10e_-6_) is indicated with a dashed horizontal line; fold change cutoff (2) is indicated with dashed vertical lines. **D** Top decreased functional categories attributed to the genes differentially expressed in diseased vs. healthy placenta EC (Ingenuity Pathway Analysis, QIAGEN). Adj. p-value <0.05. **E** Scored expression of DNA methylation writers (*DNMTs*) and editors (*TETs*) in EC of normal and diseased placenta. **F** Expression of *de novo* DNA methyltransferases in EC of normal and diseased placenta. **G** Co-staining of CD31 and DNMT3A on healthy human placenta tissue (chorionic villus zone). Scale bar 50µm.

To gain further insight into the role of EC in placenta dysfunction, differential transcriptome analysis of healthy versus diseased EC from preeclamptic placentas was performed, and revealed 359 up- and 193 down-regulated genes in preeclamptic vs. healthy EC (**Fig. 1C**). Functional annotation of these differentially expressed genes showed significant decrease of the pro-angiogenic endothelial functions “cellular movement” and “cell cycle” (**Fig. 1D**). Further analysis focused on DNA methylation regulators in diseased vs. healthy EC and revealed a down-regulation of DNA methylation writers (*DNMT3A, DNMT3B*) in EC from preeclamptic placentas, especially of *DNMT3A* (**Fig. 1E, F**). Immunohistochemical staining of healthy human placenta tissue demonstrated that DNMT3A was abundantly expressed in the endothelium of both large and small vessels in the fetal compartment (**Fig. 1G**). Fetal capillaries can be found in the chorionic villi of the human placenta. As this network of fetal capillaries is essential for gas and nutrients exchange between maternal and fetal blood, the presented data suggest a role of endothelial DNMT3A in the regulation of mature placenta function. Consequently, the reduced *DNMT3A* expression in preeclampsia EC might contribute to placenta insufficiency with compromised fetal growth.

### DNMT3A is highly expressed in fetal EC of the murine placenta labyrinth

To gain further mechanistic insight into the role of endothelial Dnmt3a in placenta function, we investigated the murine placenta labyrinth layer for analogous regulations. As in the human placenta, *Dnmt3a* was higher expressed than its homologue *Dnmt3b* (**Fig. 2A**). Furthermore, *Dnmt3a* expression increased in the course of placenta maturation, indicated by a reduction of genes involved in cell proliferation (**Fig. 2B**). In line with the prominent expression of DNMT3A in the chorionic villi of the human placenta, immunofluorescence staining analysis of the murine placenta demonstrated that DNMT3A was enriched in the fetal derived placental zones (labyrinth and junctional zone) and was not detected in the maternal derived decidua (**Fig. 2C**). Furthermore, quantification of DNA methylation via immunofluorescence staining for 5-methylcytosine (5mC) revealed increased DNA methylation levels in the labyrinth zone compared to other placental compartments, coinciding with DNMT3A expression (**Fig. 2D, 2E**). These data indicate a specific role of DNMT3A in fetal-derived zones of the placenta.

**Figure 2:**
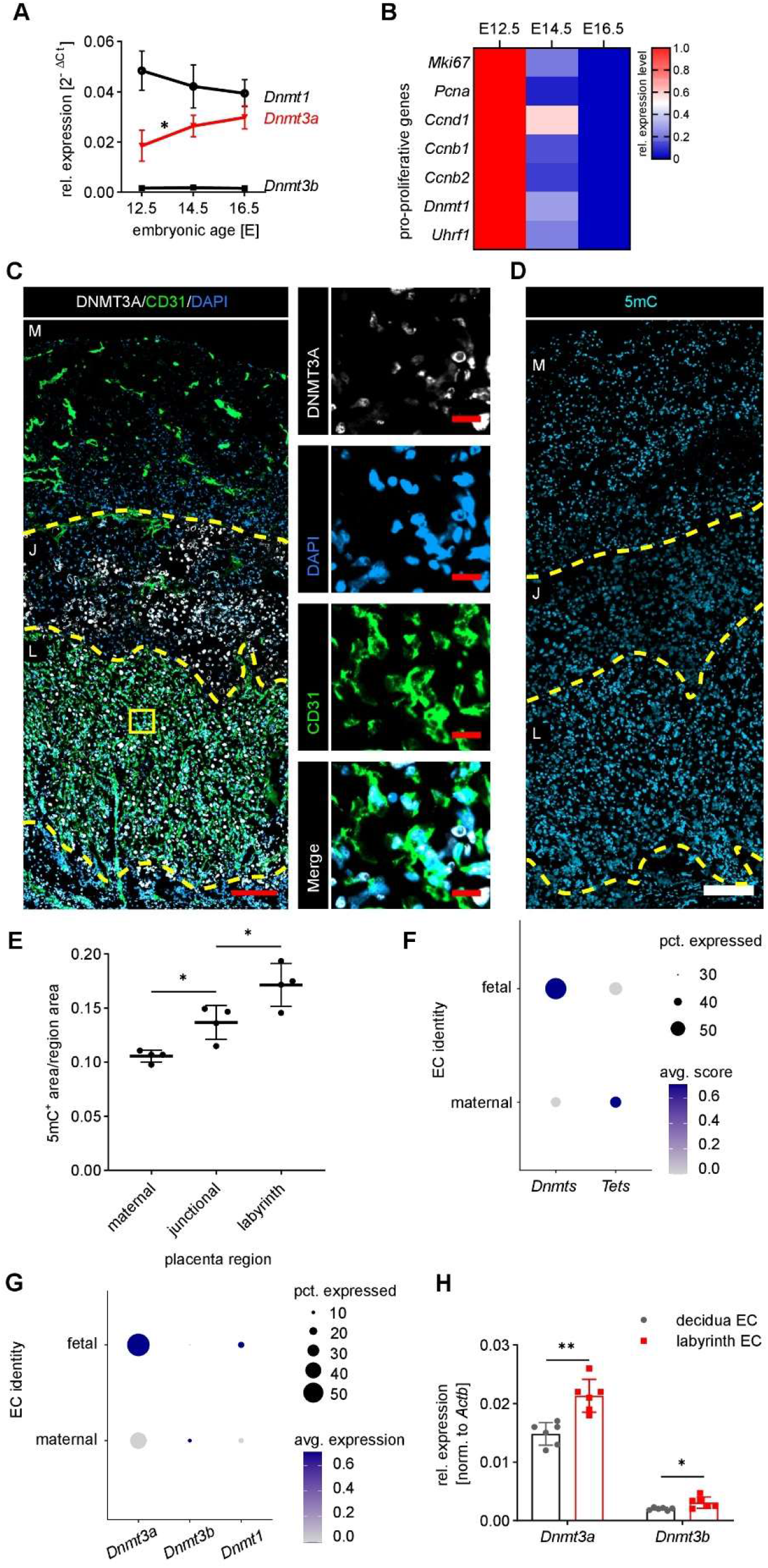
Dnmt3a is highly expressed in fetal EC in the murine placenta labyrinth. **A** *Dnmt* expression kinetics covering E12.5 to E16.5 in total mouse placenta tissue normalized to *Actb* expression. *Dnmt3a* (highlighted in red) is significantly upregulated during placenta maturation. n=5-6. **B** Heatmap depicting the expression of pro-proliferative genes in total placenta tissue. Upon placenta maturation, pro-proliferative genes are downregulated. Data is row normalized. n=5-6. **C** Co-staining of DNMT3A, CD31 (EC marker) and DAPI (nuclear marker) in the murine mature E16.5 placenta. DNMT3A is detected in the cell nucleus of EC. Placental zones are separated with yellow dashed lines. M: maternal, J: junctional; L: labyrinth. left: Scale bar 200µm. right: Zoom-in of the yellow boxed region. Scale bar 20µm. **D** Representative image of 5-methylcytosine (5mC) staining in E14.5 placenta tissue. Scale bar 200µm. **E** Quantification of 5mC intensity in different placental regions. n=4. **F** Scored expression of *Dnmts* and *Tets* in maternal and fetal EC extracted from mouse placenta single-nuclei RNA-seq (Marsh & Blelloch 2020). **G** Expression of *de novo* DNA methyltransferases in fetal and maternal EC. **H** Gene expression analysis of EC isolated from the decidua or the labyrinth of E16.5 wildtype placenta. n=6. Shown are mean±SD. Statistical significance was measured by Mann-Whitney test (A, H) or unpaired t-test (E). * p<0.05, ** p<0.01.

To investigate this finding in further detail, we split published single nuclei RNA sequencing data of mouse placenta tissue into maternal and fetal EC (see methods section for details) (Marsh and Blelloch 2020). This comparative meta-analysis revealed an elevated expression score of genes involved in the writing of DNA methylation marks in fetal vs. maternal EC (**Fig. 2F**). Specifically, *Dnmt3a* showed a higher and more abundant expression in fetal EC (**Fig. 2G**), and that difference was confirmed in sorted EC from the labyrinth (fetal) and the decidua (maternal) placental zone (**Fig. 2H**). Altogether, these murine placenta analyses underline that, similar to what we have demonstrated in human, DNMT3A is involved in the regulation of DNA methylation in the fetal endothelium of the mature placenta.

### Loss of *Dnmt3a* results in late IUGR and reduced placenta vascularization

Since the fetal endothelium within the labyrinth represents an essential exchange interface in the murine placenta, we examined the vascular phenotype and placenta function in global *Dnmt3a* knockout (KO) and wildtype (WT) embryos. Mating of mice heterozygous for the *Dnmt3a*-null allele resulted in 16% *Dnmt3a* KO mice at E16.5, representing a significant deviation from the expected Mendelian ratio which was not detectable at E10.5 (**Fig. 3A**). As the fetus switches from yolk sac to placental nutrition from day E10 onwards, this late *Dnmt3a* KO phenotype suggests a specific defect in placental function (Woods et al. 2018). Similarly, both, *Dnmt3a* KO embryo size and weight were significantly reduced at E16.5 (8.2% average reduction in size; 14.7% average reduction in weight) and not at E10.5 compared to WT littermates which indicates the late onset of IUGR upon loss of *Dnmt3a* (**Fig. 3B** - **3D**).

**Figure 3:**
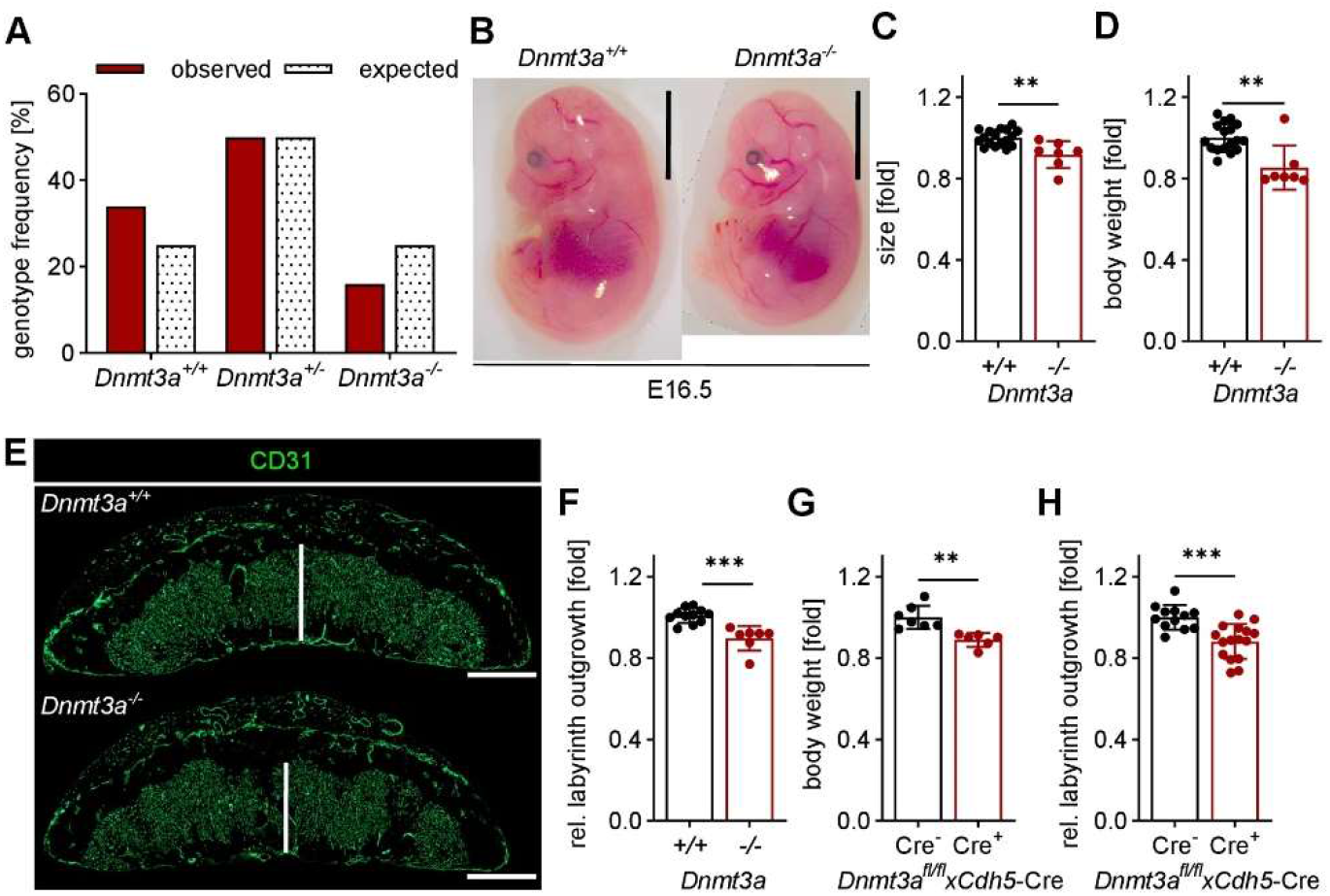
Loss of *Dnmt3a* results in late IUGR and reduced placenta vascularization. **A** Embryo genotype frequency at E16.5 from *Dnmt3a*^*+/-*^ mice crossing showing a significant deviation from the expected Mendelian ratio (Chi-square test, χ_2_=9.058, p=0.011). n=138 embryos from 17 pregnant mice. **B** Representative images of *Dnmt3a*^*+/+*^ and *Dnmt3a*^*-/-*^ E16.5 embryos. Scale bar 0.5cm. **C** Quantification of *Dnmt3a*^*+/+*^ and *Dnmt3a*^*-/-*^ embryo size at E16.5. Normalized to wildtype and per litter. n≥7. **D** Quantification of *Dnmt3a*^*+/+*^ and *Dnmt3a*^*-/-*^ embryo weight at E16.5. Normalized to wildtype and per litter. n≥7. **E** Representative images of E16.5 *Dnmt3a*^*+/+*^ and *Dnmt3a*^*-/-*^ placentas stained for the EC marker CD31. Measurement of the labyrinth outgrowth is indicated with white bars. Scale bar 1000µm. **F** Quantification of labyrinth outgrowth in *Dnmt3a*^*+/+*^ and *Dnmt3a*^*-/-*^ E16.5 placentas. n≥7. **G** Quantification of embryo weight at E16.5 of *Dnmt3a*^*fl/fl*^ mice expressing *Cdh5*-Cre (Cre+ considered as *Dnmt3a* KO in EC) or not (Cre- considered as *Dnmt3a* WT in EC). Normalized to Cre- embryos and per litter. n≥6. **H** Quantification of labyrinth outgrowth of Cre- and Cre+ *Dnmt3a*^*fl/fl*^ mice. n≥12. Shown are mean±SD. Statistical significance was measured by Mann-Whitney test (C, D, F, G, H). * p<0.05, ** p<0.01, *** p<0.001.

The labyrinth compartment of *Dnmt3a* KO placentas at E16.5 displayed a reduced vascular outgrowth, also reflected in declined CD31 (*Pecam1*, EC marker gene) expression level in total placenta tissue (**Fig. 3E, 3F**), which was not attributed to increased apoptosis in *Dnmt3a* KO placentas. Of note, in isolated *Dnmt3a*-null and WT placenta EC the expression of CD31 was unchanged. To selectively study the role of endothelial *Dnmt3a* for placenta development and function, *Dnmt3a*-floxed mice (*Dnmt3a*^*fl/fl*^) were crossed to *Cdh5*-Cre mice resulting in the loss of *Dnmt3a* in vascular EC and invasive trophoblasts (Govindasamy et al. 2021; Zhou et al. 1997). The *Cdh5*-Cre-specific loss of *Dnmt3a* partially phenocopied the effect of global *Dnmt3a* loss on embryo weight and labyrinth vascular outgrowth (**Fig. 3G, 3H**). In summary, these data suggest that *Dnmt3a* is critical for vascularization of the placenta labyrinth compartment and that its deletion causes a decreased nutrient exchange surface. The resulting limited nutrient supply to the fetal circulation then potentially accounts for fetal growth restriction of *Dnmt3a* KO fetuses.

### Loss of endothelial *Dnmt3a* reduces angiogenesis

The mouse retina represents the best characterized and most robust tool for *in vivo* angiogenesis research (Milde et al. 2013). In order to gain further insight into the reduced angiogenic potential of the *Dnmt3a* KO endothelium, retina angiogenesis in neonatal mice was analyzed. Upon loss of *Dnmt3a*, the vascularized area of the retina was significantly reduced (**Fig. 4A, 4B**). More detailed characterization demonstrated a reduction in proliferating EC and tip cells in the vascular front (**Fig. 4C - 4F**). The same phenotype was observed in a mouse model with EC-specific postnatal deletion of *Dnmt3a* via administration of tamoxifen to *Dnmt3a*^*fl/fl*^ x *Cdh5*-Cre^ERT2^ mice (*Dnmt3a*^*iECKO*^) (**Fig. 4G, 4H**). This reference model of angiogenesis strongly suggests that endothelial *Dnmt3a* is crucial for angiogenesis.

**Figure 4:**
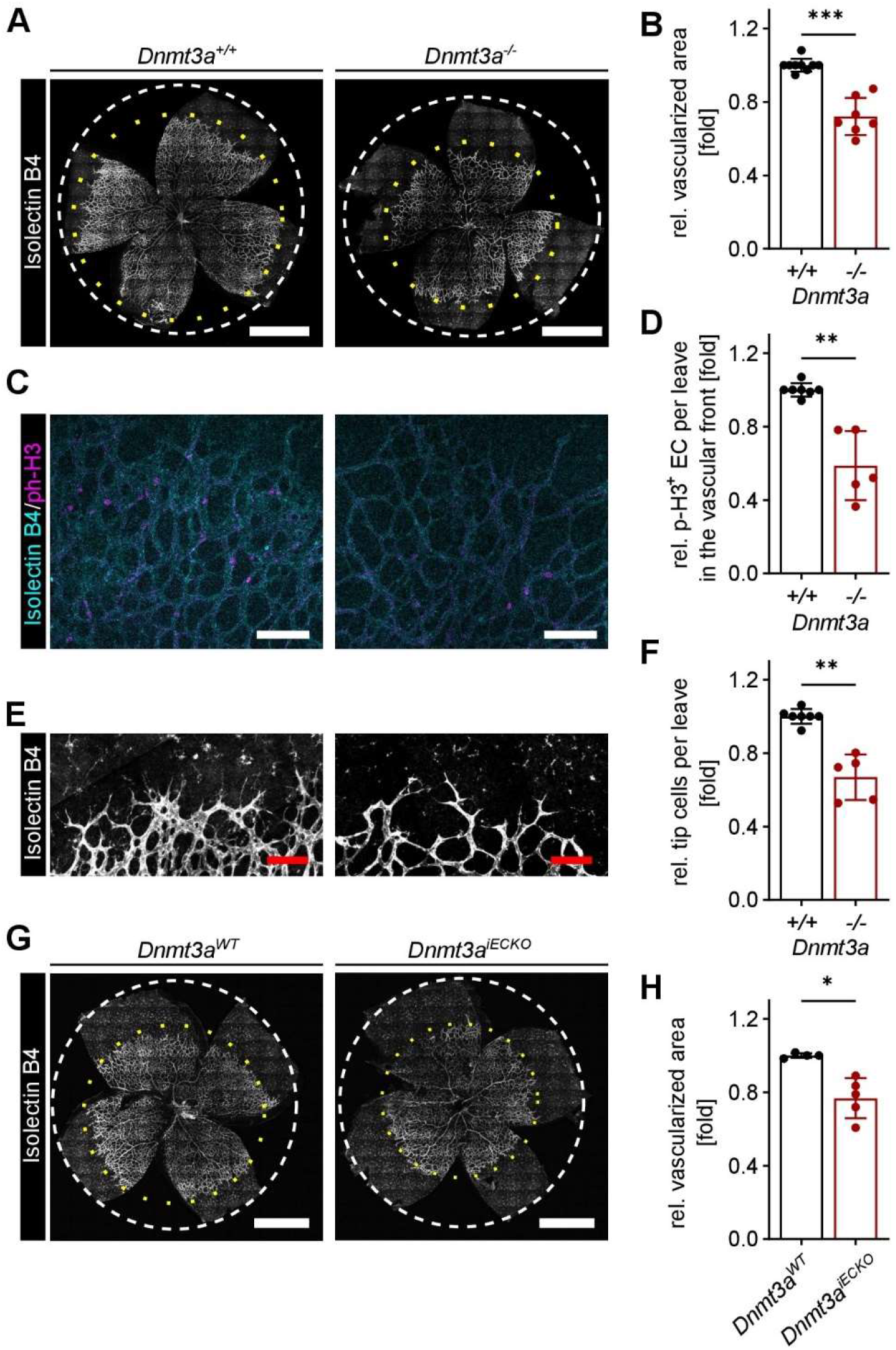
Loss of endothelial *Dnmt3a* reduces angiogenesis. **A** Representative images of postnatal day 7 (P7) retinas of *Dnmt3a*^*+/+*^ and *Dnmt3a*^*-/-*^ neonates. Vessels are stained with isolectin B4. Scale bar 1mm. **B** Quantification of the vascularized area per retina area relative to the wildtype control. n≥7. **C** Representative images of the vascular front of P7 retinas. Proliferating cells are stained by phospho-histone H3-Ser10. Scale bar 100µm. **D** Quantification of phospho-H3-Ser10-positive EC at the vascular front. Normalized to wildtype control. n≥5. **E** Representative images of the vascular front of P7 retinas. Scale bar 100µm. **F** Quantification of tip EC per leave. Normalized to wildtype control. n≥5. **G** Representative images of P7 retinas of *Dnmt3a*^*WT*^ and *Dnmt3a*^*iECKO*^ neonates. Vessels are stained with isolectin B4. Scale bar 1 mm. **H** Quantification of the vascularized area per retina area relative to wildtype control. n≥4. Shown are mean±SD. Statistical significance was measured by Mann-Whitney test (B, D, F, H). ** p<0.01, *** p<0.001.

### Loss of endothelial *Dnmt3a* results in DNA hypomethylation and loss of angiogenic capacity

Aiming at understanding the molecular mechanisms of dysfunctional angiogenesis upon loss of *Dnmt3a* in the endothelium, the DNA methylome of FACS-sorted EC (CD31^+^, CD34^+^) from postnatal day 7 *Dnmt3a* KO and WT mice was profiled by tagmentation-based whole-genome bisulfite sequencing (Weichenhan et al. 2018). Since the retina and the labyrinth zone represent poor sources for EC, pulmonary EC were isolated. The overall DNA methylation was reduced in *Dnmt3a* null EC, and 54,232 differentially methylated regions (DMRs) with a methylation difference of at least 20% were identified (**Fig. 5A**). Importantly, more than 99% of the DMRs were hypomethylated, underlining the predominant loss of DNA methylation upon loss of *Dnmt3a*. These DMRs were enriched in promoter and genic regions but underrepresented in intergenic regions compared to the reference genome (**Fig. 5B, 5C**). Region-centric functional enrichment analysis using the GREAT tool revealed a strong and significant overrepresentation of DMRs among genes involved in angiogenesis-related signaling cascades including NOTCH-, WNT-, and TGFb-signaling (**Fig. 5D**). In particular, the gene loci of the VEGF co-receptor *Nrp1* and the Notch ligand *Jag1* displayed the strongest clustering of DMRs (**Fig. 5E**).

**Figure 5:**
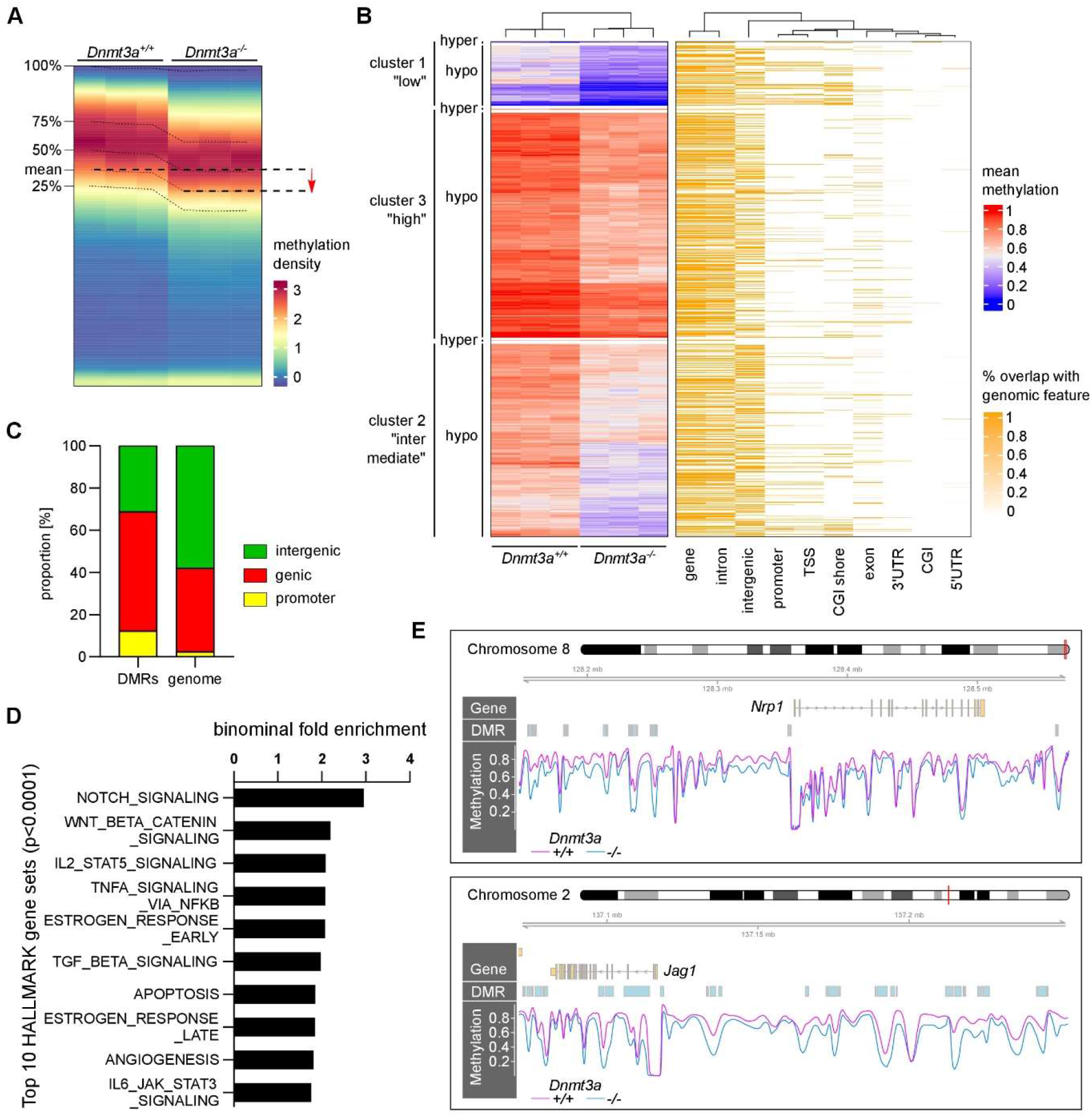
Loss of endothelial Dnmt3a results in DNA hypomethylation and loss of angiogenic capacity. **A** Global DNA methylation distribution as measured by Tagmentation-based WGBS of *Dnmt3a*^*+/+*^ and *Dnmt3a*^*-/-*^ lung EC. n=3. **B** Heat map of differentially methylated regions (DMRs) (K-means clustering, left) expanded by genomic feature annotation (right). **C** Genomic features assigned to DMRs (cutoff: 20% methylation difference) upon loss of *Dnmt3a* compared to the genomic distribution. **D** Functional enrichment analysis (GREAT) of DMRs using HALLMARK gene sets. **E** Genome browser view of gene loci encoding for *Nrp1* and *Jag1* depicting the DNA methylation level and the DMR location.

To delineate the direct effects of DNA hypomethylation on EC function from secondary effects attributed to the surrounding hypomethylated tissue, EC (HUVEC) were cultured in monoculture and DNA hypomethylation was induced by consecutive treatment with a sublethal dose of the DNA methyltransferase inhibitor decitabine (DEC). The loss of DNA methylation significantly decreased the percentage of proliferating EC, accompanied by a reduced expression of G2M-/ M-phase genes (e.g. *MKI67*) and an induction of genes mediating cell cycle arrest (e.g. *CDKN1A*). Altogether, these data suggest that the pro-angiogenic capacity of endothelial cells is compromised by genome-wide DNA hypomethylation.

### *Dnmt3a* loss is associated with a preeclampsia-like phenotype in mice

The initial characterization of *Dnmt3a* KO mouse reported a size reduction of neonatal mice followed by an early death at about one month of age, suggesting a IUGR/FGR-like phenotype (Okano et al. 1999). We also observed a reduction of born *Dnmt3a* null mice and a decreased size and weight of *Dnmt3a* KO neonates compared to their wildtype littermates. To further analyze placenta function, we focused on a potential link to placenta insufficiency in *Dnmt3a* deficient mice. Several genes that have been described to be decreased in preeclampsia in humans were downregulated in E16.5 placenta tissue of *Dnmt3a* KO compared to WT controls (**Fig. 6A**) (Cift et al. 2020; Gurtner et al. 1995; Kitazawa et al. 2017; Powers et al. 2012; Weedon-Fekjær et al. 2010). Thus, the expression profile of *Dnmt3a* null placenta tissue reflects the molecular profile of human IUGR and preeclampsia.

**Figure 6:**
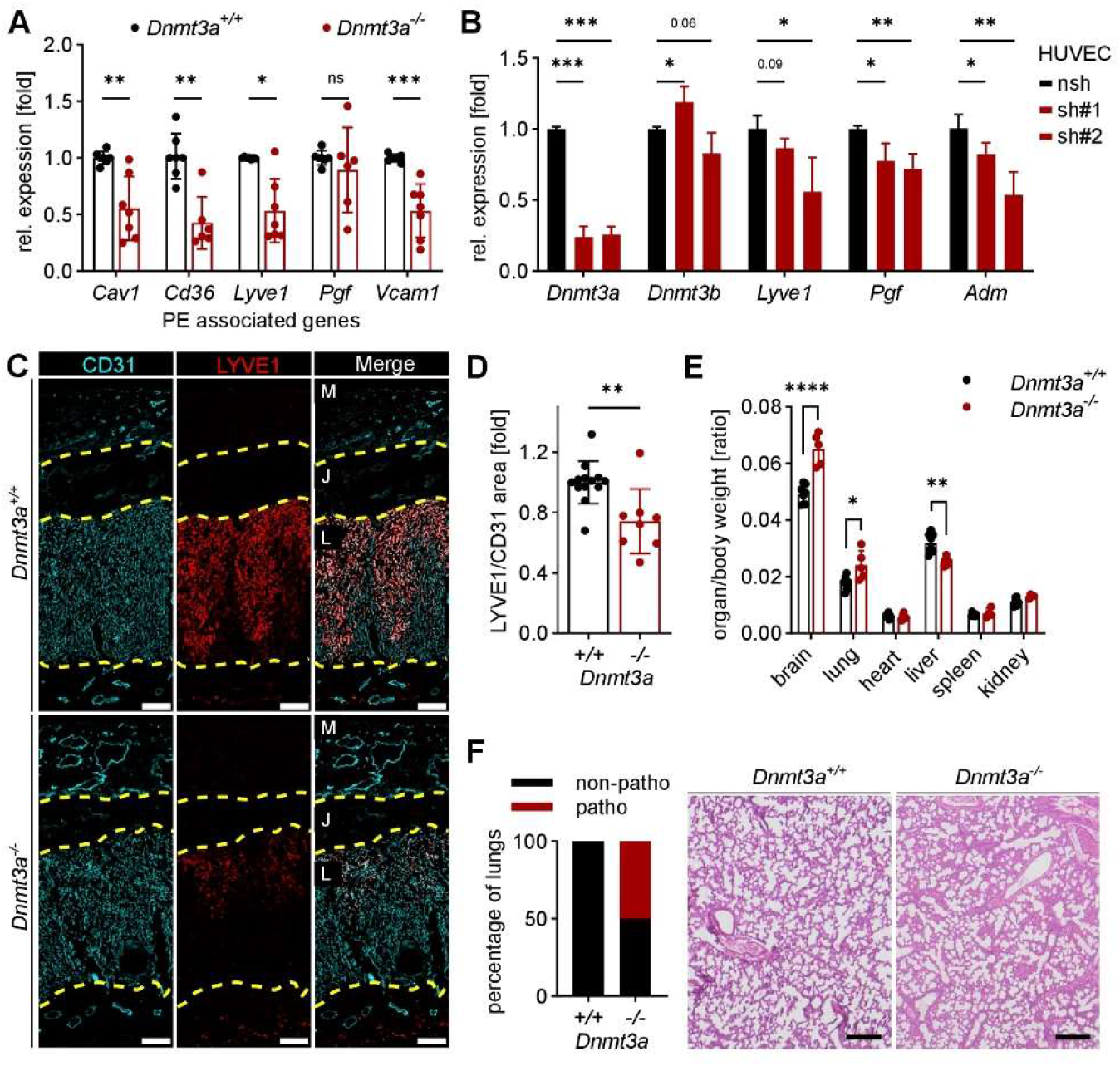
*Dnmt3a* loss is associated with a preeclampsia-like phenotype in mouse. **A** Relative expression of genes associated with preeclampsia in the murine E16.5 placenta upon loss of *Dnmt3a* vs. wildtype control. n=7. **B** Relative expression of genes associated with PE in HUVEC upon transfection with sh*Dnmt3a* (sh#1, sh#2) or control shRNA (nsh), respectively. n=3. **C** Representative images of E16.5 *Dnmt3a*^*+/+*^ and *Dnmt3a*^*-/-*^ placentas stained for CD31 and LYVE1. Scale bar 200 µm. M=maternal, J= junctional, L=labyrinth compartment. **D** Quantification of LYVE1 in the CD31-positive labyrinth area. n=8. **E** Organ to body weight ratio measured in *Dnmt3a*^*+/+*^ and *Dnmt3a*^*-/-*^ mice at P9. n≥5. **F** Analysis of lung tissue at P9 by board-certified pathologist. “non-patho” = no pathological changes, “patho” = immune infiltrates and/or fibrosis. n≥4. Shown are mean±SD. Statistical significance was measured by Mann-Whitney test (A, D, E) or unpaired t-test (B). * p<0.05, ** p<0.01, *** p<0.001, **** p<0.0001.

Many of the reported genes related to placenta dysfunction are well-known vascular genes (e.g. *Cav1, Cd36*) which are exclusively expressed in the endothelium of the placenta. To elucidate if the expression of these genes depends on the presence of *Dnmt3a* in EC, *Dnmt3a* was depleted via two independent shRNAs in cultured EC. Gene expression analysis demonstrated a significant downregulation of several genes associated with placenta insufficiency in EC but not in trophoblasts depleted for *Dnmt3a* (**Fig. 6B**). Staining of tissue sections confirmed (1) the colocalization of the vascular marker CD31 with LYVE1 and (2) the significant decrease of LYVE1 in EC (CD31-positive cells) of the labyrinth layer of the murine placenta upon *Dnmt3a* deletion (**Fig. 6C, 6D**). The reduction of LYVE1 in the labyrinth compartment was also evident after the *Cdh5*-Cre-mediated deletion of *Dnmt3a*. In contrast, VCAM1 that is not expressed by labyrinth EC but by the fetal mesenchyme, was reduced only in the global *Dnmt3a* KO mice and not in *Cdh5-Cre*-mediated *Dnmt3a* null placentas. While the global *Dnmt3a* KO mice were smaller and lighter compared to their WT littermates, *Dnmt3a*^*fl/fl*^*xCdh5*-*Cre* neonates were similar in size compared to their *Dnmt3a*^*fl/fl*^ littermates and only showed a reduced body weight. Further organ analysis in *Dnmt3a* KO neonates revealed a significantly heavier brain which indicates brain sparing, a well-known indication of nutrient deprivation as a consequence of placenta insufficiency (**Fig. 6E**) (Giussani 2016). Additionally, 50% of the lungs resected from *Dnmt3a* KO embryos showed signs of immune infiltrates and fibrosis (**Fig. 6F**). In summary, these expression analyses and postnatal evaluations suggest that the loss of *Dnmt3a* leads to vascular and thus placenta dysfunction which causes a preeclampsia-like phenotype in mice.

## Discussion

Adequate placental function is instrumental for fetal development throughout pregnancy. Placenta insufficiency is associated with fetal growth restriction – an initial manifestation of preeclampsia. The importance of adequate epigenetic regulation in placenta development, particularly in terms of the DNA methylation profile, has been shown in early studies and is recently taking center stage (Ehrlich et al. 1982; Schroeder et al. 2013; Tekola-Ayele et al. 2022). Our study took advantage of published single-cell RNA sequencing data of human (Tsang et al. 2017) and single-nuclei RNA sequencing data of mouse (Marsh and Blelloch 2020) placenta to highlight that preeclampsia is associated with reduced expression of the *de novo* DNA methyltransferase 3A in the fetal endothelium. Further phenotypic and mechanistic characterizations demonstrate that DNMT3A-dependent DNA methylation ensures the functionality of the fetoplacental endothelium for proper fetal nutrition, and loss of *Dnmt3a* leads to late placenta insufficiency in mice.

The initial characterization of *Dnmt3a* knockout mice indicated that *Dnmt3a* null embryos show a normal survival rate until E9.5 but a reduced survival after E11.5 (Okano et al. 1999). While this finding was not further addressed, our in-depth characterization of embryos, deleted for *Dnmt3a*, confirmed a normal Mendelian ratio at E10.5 and a lower ratio only later in embryonic development (E16.5). These data suggest a prominent and specific role of *Dnmt3a* late in pregnancy, which is in line with the observation that *Dnmt3a* expression is induced during placenta maturation after E12.5. Beyond these findings, we could associate the loss of *Dnmt3a* with fetal growth retardation and placenta insufficiency in addition to the described postnatally retarded phenotype in *Dnmt3a* null neonates (Okano et al. 1999). Altogether, these novel data link the *Dnmt3a* null phenotype in mice with placental dysfunction.

Proper DNA methylation is essential in placenta and fetal development. Upon fertilization, the initial global demethylation is followed by *de novo* DNA methylation predominantly in cells with fetal fate resulting in global hypermethylation of fetal cells compared to other placental cells (Guo et al. 2014; Smith et al. 2014; Zhang et al. 2021; Santos et al. 2002). Based on these findings, fetal cells in the placenta are likely the cell population being most susceptible to wide-spread loss of *de novo* DNA methylation. A reduction of *Dnmt3a* in fetal capillaries was associated with fetal growth arrest (Gu et al. 2017). Along the same line, we show that the fetoplacental endothelium is expressing high levels of DNMT3A in healthy placentas and the loss of DNMT3A-dependent DNA methylation in the endothelium results in decreased labyrinth vascularization and fetal growth defects. Thus, several lines of evidence suggest that *de novo* DNA methylation in the fetal endothelium plays a crucial role in placental maturation and thereby contributes to proper fetal nourishment.

Based on this notion, our data imply that dysfunctional fetoplacental vasculature is central in FGR cases and preeclampsia. In fact, it has been described that the size of the labyrinth zone determines its nourishment capacity underlining its critical role in the pathogenesis of FGR (Woods et al. 2018). Many studies have reported a dysregulation of endothelial-specific genes that are now used as predictive markers of preeclampsia (MacDonald et al. 2022). For instance, aberrant levels of circulating placental growth factor (PGF) and soluble fms-like tyrosine kinase 1 (sFlt1) are consistently identified in preeclampsia and, therefore, used as test-candidates in the clinics. Pilot studies have focused on the removal of anti-angiogenic proteins with beneficial effects (Thadhani et al. 2011, 2016). In summary, the vascular link of preeclampsia is well established and this present study provides specific evidence for the central importance of the fetoplacental circulatory system for appropriate placental function and fetal development.

Several studies report that aberrant DNA methylation is associated with placenta-related pathologies (Cirkovic et al. 2020; Nelissen et al. 2011; Wilson et al. 2018), and few studies link a reduction of DNMT3A with placental insufficiency (Jia et al. 2017; Gu et al. 2017). Yet, the underlying cause of DNA methylation loss or DNMT3A impairment in late pregnancy remains unknown. On the one hand, dysregulation of epigenetic candidate molecules that cooperate with DNMT3A could compromise DNMT3A function. For instance, in cancer UHRF1 represents a key epigenetic regulator required for DNMT3A-dependent DNA methylation (Jia et al. 2016). In the same line, the epigenome of embryonic stem cells is dependent on the interplay of active DNA demethylation via TET (Ten-eleven translocation) proteins and re-methylation by DNMTs (Gu et al. 2018). On the other hand, global placental DNA hypomethylation has also been associated to air pollution and Pb exposure (Janssen et al. 2013; Tasin et al. 2022). These studies demonstrate the potentially far-reaching consequences of the dysregulation of single molecules or broad environmental exposures for adequate placental function and fetal nutrition.

In summary, this study reveals that endothelial DNMT3A represents a key epigenetic regulator in the placenta. By the establishment of an appropriate DNA methylation landscape, DNMT3A is required for maintaining the angiogenic capacity of EC and consequently appropriate tissue vascularization. Finally, *Dnmt3a* loss in the endothelium can be associated with placenta insufficiency resulting in reduced fetal growth in humans and mice. Thus, our study provides valuable insight into the role of the fetoplacental vasculature in general and *Dnmt3a* especially in controlling placenta function in health and disease.

## Methods

### Human single-cell data analysis

The human placenta single-cell RNA sequencing data was provided on request from Tsang et al. 2017. Data was analyzed in R (version 4.0.2) using the package Seurat (version 4.0.2). Cells with fewer than 200 or greater than 2500 unique genes and with greater than 5% mitochondrial counts, were excluded from analyses. For the analysis of normal placenta cells, individual healthy placenta samples were merged and counts were scaled and log2-normalized using ScaleData and NormalizeData, respectively. PCA was performed with the 2000 most highly variable features (assessed with FindVariableFeatures as input). Significant dimensions were assessed using ElbowPlot and clustering was performed using 15 dimensions and a resolution of 0.3 with FindNeighbors and FindClusters, respectively. Genes for cluster annotation were extracted from Tsang et al. 2017 and visualized with FeaturePlot. To assess the log2-normalized expression of epigenetic modifiers in the defined cell populations, gene lists for “chromatin remodeling”, “histone modification” and “DNAme alteration” were defined (Plass et al. 2013). The “DNAme alteration” signature was further separated into “writer”, including *DNMT1, DNMT3A* and *DNMT3B*, and “editor” including *TET1, TET2* and *TET3*. The expression scores for the defined gene lists were added using AddModuleScore and data was visualized with DotPlot. The expression of individual genes was as well generated with the dotplot function. To extract EC from healthy and diseased placentas, an EC score was assigned to each cell based on the expression of manually curated key EC genes (*CD34, CDH5, ERG, TIE1, TEK* and *VWF*) using AddModuleScore. Cells with an EC score higher 0.4 were selected as a subset for further analysis. EC from healthy and diseased placentas were merged with Merge. Clustering was performed as described above with a dimension of 9 and a resolution of 0.5. DimPlot was used to show the EC score expression in the extracted EC and their health state. The differential gene expression analysis in extracted EC was performed based on their health state (healthy vs. diseased) using the Wilcoxon Rank Sum test. Differentially expressed genes were assessed with FindAllMarkers. Only genes that were detected in a minimum fraction of 10% of cells and passed the threshold of 0.25fold change were used. Data was visualized as volcano plot with EnhancedVolcano (version 1.8.0). Functional gene annotation was performed through the use of QIAGEN Ingenuity Pathway Analysis (Krämer et al. 2014).

### Mouse single-cell data analysis

The mouse placenta Seurat object was obtained from Marsh and Blelloch 2020. The package Seurat (version 4.0.2) was used for data analysis. Cells with fewer than 600 or greater than 3000 unique genes and with greater than 12% mitochondrial genes were excluded from the following analysis. PCA was performed using RunPCA with the 2000 most highly variable features (assessed with FindVariableFeatures) as input. Clustering was performed using 17 dimensions and a resolution of 0.5 with FindNeighbors and FindClusters, respectively. To extract EC, an manually curated EC score was assigned to each cell based on the expression of key EC genes (*Pecam1, Erg, Tie1, Tek* and *Kdr*). Cells with an EC score higher 0.75 were used for further analysis. The data was log2-normalized and scaled with NormalizeData and ScaleData, respectively and further processed as described above. For cell clustering 13 dimensions with a resolution of 0.5 were used. To distinguish between maternal and fetal EC, that dataset was reduced to male placenta samples (E12.5 and E14.5). Discrimination of maternal and fetal EC was assessed based on *Xist* expression. The dataset contained EC with either low (n=890, male) or high (n=58, female) *Xist* expression. The difference in the number of gender-specific cells was due to the selective enrichment for the labyrinth layer. The expression of epigenetic modifiers was assessed as described above.

### Mice

Mice were housed in individually ventilated cages under pathogen-free conditions. Animals had free access to food and water and were kept in a 12h light-dark cycle. All mice were handled according to the guidance of the Institute and as approved by the German Cancer Research Center. Heterozygous *Dnmt3a* knockout mice were obtained from Jackson Laboratory (# No. 018838) to generate homozygous *Dnmt3a* knockout mice (approved by the Competent Authorities in Karlsruhe, Germany; permit G-82/19). C57BL/6 *Dnmt3a*^*flox/flox*^ mice (RIKEN BioResource Center, No. RBRC03731) were crossed with C57BL/6 *Cdh5-Cre* mice (*B6;129-Tg(Cdh5-cre)1Spe/J;* Jackson Laboratory). For postnatal *Dnmt3a* deletion, C57BL/6 *Dnmt3a*^*flox/flox*^ mice were crossed with C57BL/6 *Cdh5-Cre*^*ERT2*^ mice (*Tg(Cdh5-cre/ERT2)1Rha*; kindly provided by Dr. Ralf Adams, MPI Münster, Germany) to specifically delete *Dnmt3a* in *Cdh5*-expressing cells upon 4-Hydroxytamoxifen application (permit: G-283/18). Cre-negative mice, equally treated with 4-Hydroxytamoxifen, were used as controls in all experiments. Pups were intragastrically injected with three doses of 4-Hydroxytamoxifen (50µg) (Sigma, 68047-06-3) at P2, P3 and P5 dissolved in ethanol and peanut oil.

### Human sample collection

Human full term placenta samples that were considered having a normal placentation, were collected immediately after resection. Tissue was dissected from the fetal side of the placenta and embedded in O.C.T. Human material collection was approved by the Competent Authorities in Tübingen, Germany (S-320/2022).

### EC isolation by FACS

To sort maternal and fetal placenta EC, the isolated placentas were cut in the sagittal plane. From each half, the decidua was removed with a micro-scissor without disturbing the labyrinth. Labyrinth and decidua samples were digested with 2% DNAseI (ROCHE)/Liberase (Liberase™ TM Research Grade, MERCK, 6.5U) in DMEM (GIBCO) for 15min at 37°C. After the tissue was gently disintegrated by passing the mixture through an 18-G needle, the samples were incubated for another 15min at 37°C. The cell suspension was passed through a 100μm cell strainer. Erythrocyte lysis was performed by resuspending the cell pellet in ACK buffer following an incubation for 5min at RT. To block non-specific binding, samples were incubated with anti-mouse CD16/32 antibody (Mouse BD Fc Block™, BD Pharmingen™). Cells were washed and incubated with the antibody staining mix, before sorting the samples. Dead cells were excluded by propidiumiodid (PI) staining. Live CD45^-^/CD362^-^/Pdpn^-^/Ter119^-^/CD31^+^ cells were sorted with a BD FACSAria ll (BD Biosciences, Heidelberg, Germany). Isolation of mouse lung EC was performed as described in (Schlereth et al. 2018).

### Tagmentation-based whole genome bisulfite sequencing (T-WGBS)

T- WGBS was performed from FACS-sorted EC (CD31^+^, CD34^+^) from postnatal day 7 *Dnmt3a* KO and WT mice in triplicates as described in (Schlereth et al. 2018). Differentially methylated regions (DMRs) were detected by three different tools separately: the Bioconductor package bsseq (Hansen et al. 2012) (version 1.28.0, cutoff was set to |t-value| > 4.6), the Bioconductor package DSS (Park and Wu 2016) (2.40.0, cutoff was set to p.threshod < 0.01), and the command-line tool metiline (Jühling et al. 2015) (version 0.2-7, cutoff was set to q-value < 0.05). The union of the three DMR lists was used as the final set of DMRs. DMRs were assigned to the nearest gene TSS. Functional enrichment analysis on DMRs was applied with the GREAT method (McLean et al. 2010) with the Bioconductor package rGREAT version 1.99.3 (Gu and Hübschmann 2022). Hallmark gene sets for mouse were obtained by the R package msigdbr (version 7.5.1). The R package gviz was used to generate locus plots (Hahne and Ivanek 2016). The methylome data have been deposited at the NCBI Gene Expression Omnibus (GEO) and will be made available upon request.

### RNA isolation, reverse transcription and quantitative measurement of RNA

Snap frozen tissue was mechanically disrupted with 5mm sized metal beads in TRI Reagent® (Sigma, T9424), following RNA extraction using phenol:chloroform. RNA was purified using the GenElute Mammalian total RNA Miniprep kit (MERCK). Samples from cell culture were harvested in the RNA extraction buffer supplemented with 10mM ß-ME. RNA was purified using the GenElute Mammalian total RNA Miniprep kit according to the manufacturers’ recommendation.

Reverse transcription was performed using the QuantiTect® Reverse Transcription Kit (QIAGEN) according to the manufacturer’s instructions. TaqMan™ Fast Advanced Mastermix (ThermoFisher Scientific) was used to detect differences of mRNA transcription levels. Reactions were performed in 384-well format on the StepOnePlus Real-Time PCR system.

### Retina analysis

Postnatal day 7 pups were used for retina angiogenesis analysis. The retina vasculature develops in the first week after birth radially from the optic nerve to the periphery. The eyeball was fixed in ice-cold methanol at -20°C. Blocking and permeabilization of the isolated retina were performed with 10% normal goat serum/ 0.5% Triton-X 100/ 1% BSA in PBS. The retinal vasculature was stained by incubation with isolectin B4 and primary antibody diluted in 0.2% Tween-20/ 1% BSA in PBS, followed by incubation with appropriate secondary antibodies. Fluorescently labeled retinas were flat-mounted with DAKO mounting medium on microscope slides and imaged using the confocal Zeiss LSM710 microscope with a 20x/ 0.4 Dry objective. Image analysis was done in ImageJ.

### Staining of tissue sections and analysis

For immunofluorescent staining, Tissue-Tek O.C.T. embedded cryopreserved tissue was cut into 7µm sections and fixed in ice-cold methanol. For immunohistochemical stainings, paraffin embedded tissue was cut and sections were rehydrated, following antigen retrieval with citrate buffer. Blocking and tissue permeabilization was performed in 10% normal donkey serum/ 3% BSA, followed by incubation with the appropriate antibodies. DAPI was used for nuclei visualization. Slides were mounted using DAKO mounting medium and images were taken with the Zeiss Axio Scan.Z1 using an air 20×/ 0.8 Plan-APOCHROMAT objective. Image analysis was done in ImageJ.

### Cell culture

HUVEC were purchased from PromoCell and cultured in Endopan 3 supplemented with 3% FCS and supplements (PAN Biotech) at 37°C, 5% CO_2_ and high humidity maximum until passage 6. Cell culture plates were coated with 0.1% gelatin prior to cell seeding. Prior to cell proliferation analysis, cells were starved in medium without supplements for 1h. EdU was added at a final concentration of 10µM for 3h. Harvesting, fixation, permeabilization and staining were performed using the Click-iT EdU Flow Cytometry Assay Kit Alexa Fluor 488 (ThermoFisher Scientific) according to the manufacturer’s protocol. Annexin V assay was performed using the FITC conjugated EBioscience™ Annexin V Apoptosis Detection Kit (Invitrogen™) according to the manufacturer’s protocol. FxCycle Violet stain was used instead of PI. Cells were analyzed on a BD FACSCanto II (BD Bioscience). For DNA de-methylation, HUVEC were treated with 500nM 5-Aza-2’-deoxycytidine (MERCK) dissolved in 0.9% NaCl. Cells were passaged two times before harvesting. The media was changed every day. JEG3 (HTB-36) were purchased from ATCC and cultured in EMEM supplemented with 2 mM L-glutamine, 1% Non-Essential Amino Acids, 1mM sodium pyruvate (CLS), 10% FCS and 1% Penicillin Streptomycin Solution at 37°C, 5% CO_2_ and high humidity. Upon lentiviral transduction with pGIPZ-shRNA encoding virus, cells were selected with puromycin (0.4 µg/ml) for 4-6 days.

### DNA methylation analysis via dot blot

Genomic DNA was isolated using the NucleoSpin Tissue – Mini kit for DNA from cells and tissue (MACHEREY-NAGEL). Genomic DNA was sheared with a 30-G syringe. DNA was mixed with 20µl denaturation buffer (1M NaOH, 25mM EDTA) and boiled for 10min at 95°C. Samples were placed on ice and mixed with 50µl neutralization buffer (2M ammonium acetate, pH 7.0). Nitrocellulose membranes were fixed in a microfiltration blotting device (Bio-Dot Apparatus). DNA was applied and membranes were washed and air-dried following UV-crosslinking. After blocking with 5% milk in TBS-T for 1h at RT, incubation with primary antibodies was performed at 4°C overnight. After washing, the HRP-conjugated secondary antibody was applied for 1h at RT. After washing, membranes were developed with ECL solution (Thermo Scientific™ Pierce™).

### Statistical analysis

Statistical analyses were performed using GraphPad Prism version 8. Data are expressed as mean ± SD. A p-value of <0.05 was considered statistically significant and marked by asterisks (* p<0.05, ** p<0.01, *** p<0.001). Unless otherwise stated, n represents the number of independent mice or samples in biological replicates analyzed per group or condition.

## Acknowledgments

We would like to acknowledge the excellent technical support of Claudine Fricke, Luisa Knospe and Benjamin Schieb (H.G.A. laboratory) as well as Marion Bähr (C.P. laboratory). We thank the Flow Cytometry Core Facility and the Central Animal Laboratory, German Cancer Research Center (DKFZ), for providing excellent services. We thank the High Throughput Sequencing unit of the Genomics and Proteomics Core Facility (DKFZ) for providing excellent sequencing and QC analysis services. We thank the Light Microscopy Core Facility (DKFZ) for providing instruments. This work was supported by a grant from the Deutsche Forschungsgemeinschaft (DFG) (project A5 within CRC1366 “Vascular Control of Organ Function” [project number 39404578]) to KS and the German-Israeli Helmholtz International Research School Cancer-TRAX (HIRS-0003) to GS.

## Author contributions

S.G. and K.S. conceived the study. G.S., E.G., D.W., and C.M. performed the experiments. G.S., M.J., Z.G., and K.S. analyzed sequencing data. G.S. and K.S. wrote the original draft of the manuscript. M.J., E.G., Z.G., D.W., C.M., M.S., C.P., and H.G.A. reviewed and edited the manuscript. K.S. and H.G.A supervised the study, and acquired the funding.

